# Enrichment of specific GABAergic neuronal types in the mouse perirhinal cortex

**DOI:** 10.1101/2022.01.30.478360

**Authors:** Maximiliano José Nigro, Kasper Kjelsberg, Laura Convertino, Rajeevkumar Raveendran Nair, Menno P. Witter

## Abstract

GABAergic neurons represent 10-15% of the neuronal population of the cortex but exert a powerful control over information flow in cortical circuits. GABAergic neurons show an extraordinary diversity in their morphology, physiology, molecular markers and connectivity. This diversity allows GABAergic neurons to participate in a wide variety of microcircuit motifs. The diversity of GABAergic neurons has been shown to be conserved across cortical regions. The GABAergic population can be broadly divided in three major classes parvalbumin, somatostatin and 5HT3aR groups. The largest GABAergic class in the cortex is represented by the parvalbumin-expressing fast-spiking neurons, which provide powerful somatic inhibition to their postsynaptic targets. Recently, the density of parvalbumin-expressing neurons has been shown to be lower in associative areas of the mouse cortex, including the perirhinal cortex, as compared to sensory and motor areas. In the present study we investigated whether this reduction in parvalbumin-expressing neurons leads to a decreased GABAergic population, or to an enrichment of other GABAergic cell-types. We found that the GABAergic population of the perirhinal cortex is comparable to that of a primary sensory area, and it is enriched of neurons belonging to the 5HT3aR group. We also demonstrate that, despite the low density of parvalbumin-expressing neurons, the perirhinal cortex contains a comparable population of fast-spiking neurons, most of which do not express parvalbumin. Our results demonstrate a yet uncharacterized diversity within the fast-spiking population across cortical regions.

## Introduction

The wide neuronal diversity of the cerebral cortex is thought to endow it with a variety of circuit motifs that shape information processing (Harris and Shepherd 2015; Luo 2021). The cerebral cortex can be parcellated in functional areas that show specific cytoarchitectonic features and molecular marker expression (van Essen and Glasser, 2018). The relationship between cortical parcellation and distribution of neuron types in the cortex is poorly understood. GABAergic neurons represent 10-15% of the neuronal population and their diversity allows them to participate in different cortical microcircuits (Fishell and Kepecs, 2014; Tremblay et al., 2021). GABAergic neurons are divided in two major groups according to embryonal origin: those derived from the medial ganglionic eminence (MGE) and those derived from the caudal ganglionic eminence (CGE) (Rudy et al 2011). MGE-derived GABAergic neurons are further divided according to molecular marker expression into parvalbumin (PV)-expressing and somatostatin (SST)-expressing interneurons (INs). CGE-derived interneurons (CGE-INs) are labeled by GFP in the 5HT3aR-GFP mouse line (Lee et al 2010). Transcriptomic analysis has confirmed this classification and further divided the CGE group into three major classes: vasoactive intestinal peptide (VIP)-expressing, Lamp5, and Sncg groups (Tasic et al 2018). Molecularly defined GABAergic neurons have been shown to be homogeneously distributed across sensory-motor areas of the neocortex (Xu and Callaway 2010). However, a recent cortex-wide examination of the distribution of three major GABAergic groups (PV, SST, VIP) revealed regional specializations (Whissell et al 2015; Kim et al 2017). In particular, the density of PV-INs decreases in association areas, including the perirhinal cortex (PER). PV-INs are the most abundant GABAergic type in the neocortex and provide powerful perisomatic inhibition to their postsynaptic targets (Tremblay et al., 2016). 79 PER is part of the parahippocampal region and represents a gateway for sensory information entering the hippocampal formation (Witter et al., 2000), by excerting an inhibitory control on the information travelling into the hippocampal network (DeCurtis and Paré, 2004). The mechanisms underlying this inhibitory control are still poorly understood. One possibility is that this inhibitory gating relies on strong perisomatic feedforward inhibition mediated by PV-INs (Willems et al 2018). This is in striking contrast with the low density of PV-INs in PER. The low density of PV-INs in PER suggests that either the GABAergic population of PER is smaller than that of the neocortex, or that other GABAergic cell-types are enriched. In the present study we aimed to explore these two scenarios by combining mouse genetics, molecular marker expression, enhancer mediated transgene expression and electrophysiological characterization of GABAergic neurons in PER. We compared the GABAergic population of PER to that of the area of whisker representation of the primary somatosensory cortex (wS1), whose GABAergic diversity has been extensively described (Lee et al., 2010; Feldmeyer et al., 2018). We found that indeed the GABAergic population is not smaller in PER and contains a higher density of 5HT3aR-expressing interneurons. Characterization of the firing patterns of GABAergic neurons in PER shows that this is partially due to an enrichment of late-spiking neurogliaform cells. Most strikingly, we also found that the percentage of fast-spiking neurons was much higher than expected from the density of PV-INs. Using an enhancer-driven viral approach, we demonstrate that the population of fast-spiking neurons in PER includes PV-expressing neurons and a population of neurons not expressing PV.

## Results

### Comparison of the GABAergic population of wS1 and PER

To compare the whole GABAergic population of PER and wS1 we used the GAD67-EGFP mouse line and performed immunofluorescence staining for PV, SST and GFP (Figure 1A-D). We quantified the number of PV-IR, SST-IR and GFP-IR/PV-SST- (neurons labeled by the GAD67-EGFP mouse line but not expressing either PV or SST) neurons across layers of the two cortical areas and calculated their densities. We performed a PCA on the densities of these markers across the layers of wS1 and PER (A35 and A36) to test how these cell-types describe these two cortical regions (Figure 1E). The first two principal components accounted for 93.8% of the variance and the data was well segregated in the principal component space (Figure 1E and Figure 1 supplement 2A). The variables that best explained the variance were the density of PV-INs in layers 2/3-6, which was lower in PER, and the density of GFP-IR/PV-SST-in layers 2/3-6, which was higher in PER (Figure 1E and Figure 1 – figure supplement 2B). The PCA results were confirmed by a statistical comparison of the three molecularly defined GABAergic populations (Figure 1 – Figure supplement 3). Indeed, the density of PV-INs was significantly lower in layer 2/3-6 of PER, whereas the density of GFP/-PV-SST neurons was significantly higher in L2/3-6 as compared to wS1 (Figure 1 – Figure supplement 3). The PCA analysis showed a slight contribution of SST-INs in L2/3, and indeed their density was significantly higher in L2/3 and 6 of PER (Figure 1E and Figure 1 – Figure supplement 3). Our data show that A35 and A36 are very similar in their GABAergic population. However, the densities of these GABAergic types in A36 are between that of A35 and wS1 (Figure 1E and F, Figure 1 – Figure supplement 3). These results confirmed previous reports of low density of PV-INs in PER (Whissel et al., 2015; Kim et al., 2017), and further show that PER is enriched in a population of GABAergic neurons that does not express either PV or SST.

**Figure 1.**
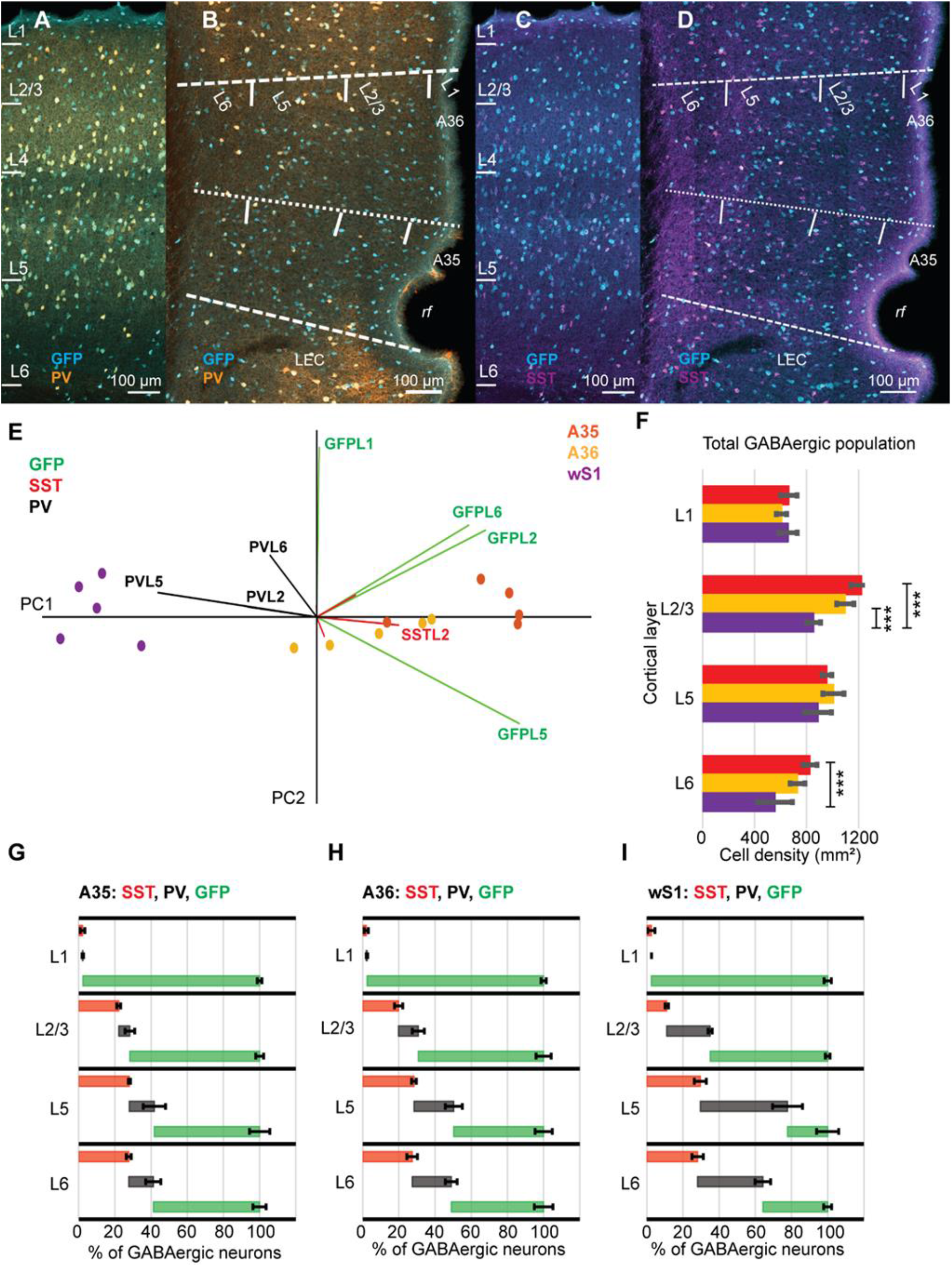
**A**. Representative confocal stack of immunofluorescence for GFP (turquoise) and PV (orange) in the wS1 of a GAD67-GFP mouse. **B**. Representative confocal stack of immunofluorescence for GFP and PV in PER of a GAD67-GFP mouse. **C**. Representative confocal stack of immunofluorescence for GFP (turquoise) and SST (orange) in the wS1 of a GAD67-GFP mouse. **D**. Representative confocal stack of immunofluorescence for GFP and SST in PER of a GAD67-GFP mouse. **E**. Graph plotting the samples in the principal component space using PC1 and PC2. **F**. Bar plot showing the statistical comparison of the density of GABAergic neurons across the layers of A35 (red), A36 (yellow) and wS1 (purple) (n= 5 mice): L1 (F= 1.33, df=2, p= 0.29, One-Way ANOVA); L2/3 (F= 37.4, df= 2, p= 6.9E-6, One-Way ANOVA; A35 vs wS1, p= 7.6E-6, Bonferroni correction; A36 vs wS1, p= 0.0002, Bonferroni correction); L5 (F= 2.8, df= 2, p= 0.1, One-way ANOVA); L6 (F= 11.5, df= 2, p= 0.002, One-Way ANOVA; A35 vs wS1, p= 0.008, Bonferroni correction). **G**. Cumulative percentage of SST-IR (red), PV-IR (black) and GFP-IR/-PV-SST neurons in the total GABAergic population of A35. **H**. Cumulative percentage of SST-IR (red), PV-IR (black) and GFP-IR/PV-SST-neurons in the total GABAergic population of A36. **I**. Cumulative percentage of SST-IR (red), PV-IR (black) and GFP-IR/PV-SST-neurons in the total GABAergic population of wS1.

Finally, we show that PER has a higher density of GABAergic neurons, particularly in L2/3 and L6 (Figure 1F). Our results suggest that in PER the low density of PV-INs is compensated by an enrichment of other classes of GABAergic neurons. A comparison of the percentage of marker-IR neurons in the GABAergic population further confirmed the reduction of PV-INs and an increased representation of GFP-IR/-PV-SST neurons across the layers of A35 and A36 (Figure 1G-I).

### PER is enriched in GABAergic cell-types of the 5HT3aR group

In sensory and motor areas of the dorsal cortex, PV-INs represent the largest group of GABAergic neurons (Tremblay et al., 2016). Our results show that in PER the largest population across layers does not express either PV or SST (Figure 1). We reasoned that this population might be in part composed of neurons expressing GFP in the 5HT3aR-EGFP mouse line, which do not express either PV or SST (Lee et al., 2010). We tested this hypothesis by quantifying the density of GFP labeled neurons in the 5HT3aR-EGFP mouse line (Gong et al., 2003) (Figure 2). We found that PER contains a significantly higher density of GFP labeled neurons as compared to wS1, particularly in L2/3, 5 and 6 (Figure 2C). We then tested whether VIP-INs contributed to this difference by performing immunolabelling for VIP in the GAD67-EGFP mouse line (Figure 2 – Figure supplement 1). The density of VIP-INs in PER tended to be higher than in wS1, but the difference was not significant, other than in L5 (Figure 2 – Figure supplement 1). We conclude that the VIP-INs are not the major source of the enrichment of 5HT3AR-INs in PER.

**Figure 2.**
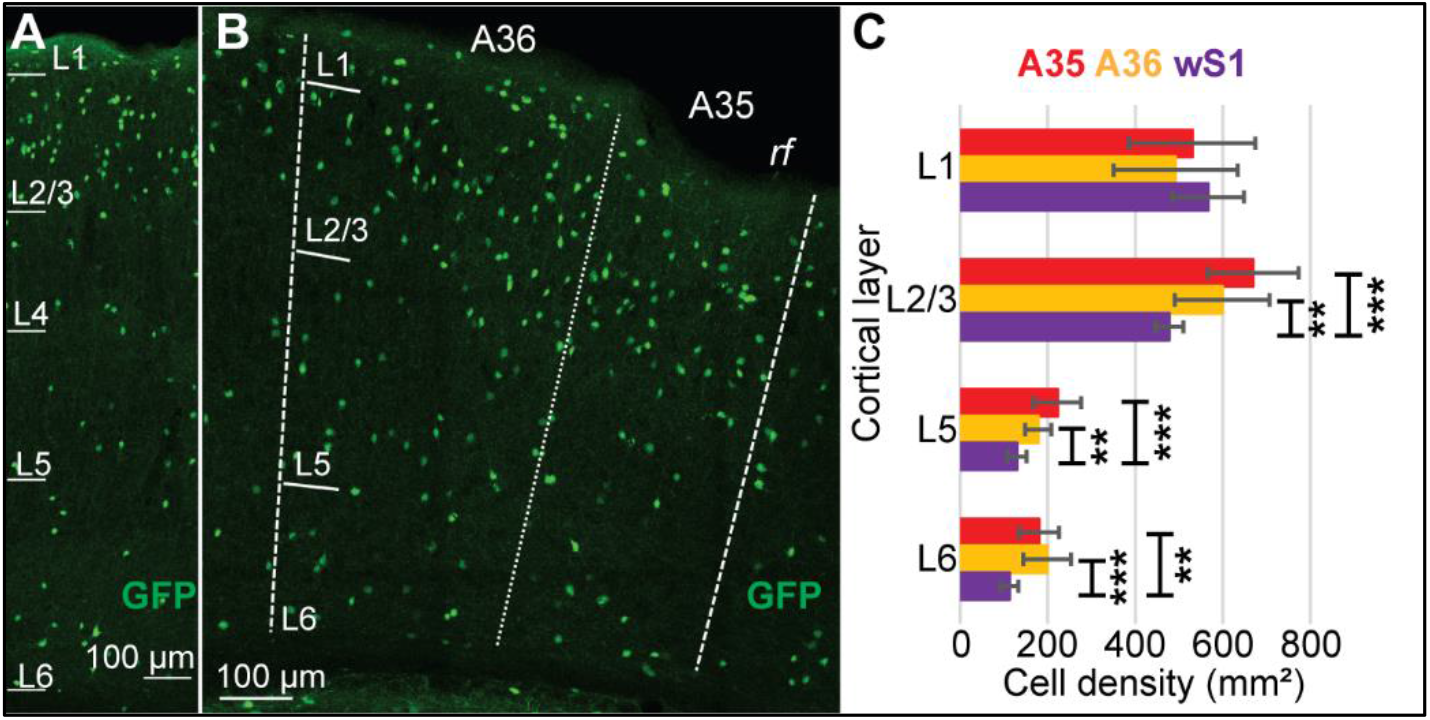
**A**. Representative confocal stack of immunofluorescence for GFP (green) in wS1 of the 5HT3aR-EGFP mouse line. **B**. Representative confocal stack of immunofluorescence for GFP (green) in PER of the 5HT3aR-EGFP mouse line. **C**. Bar plot showing the statistical comparison of the density of GFP-IR neurons in the 5HT3aR-EGFP mouse line (n= 10 slices in each area): L1 (F= 0.78, df= 2, p= 0.47, One-way ANOVA); L2/3 (F= 10.81, df= 2, p= 0.0004, One-way ANOVA; A35 vs wS1, p= 0.0003, Bonferroni correction; A36 vs wS1, p= 0.009, Bonferroni correction); L5 (F= 13.24, df= 2, p= 9.84E-5, One-way ANOVA; A35 vs wS1, p= 0.0005, Bonferroni correction; A36 vs wS1, p= 0.001, Bonferroni correction); L6 (F= 10.02, df= 2, p= 0.0006, One-way ANOVA; A35 vs wS1, p= 0.002, Bonferroni correction; A36 vs wS1, p= 0.0009, Bonferroni correction).

### L5 of PER is enriched of late-spiking neurogliaform neurons

The 5HT3AR-INs group includes VIP-INs, Lamp5-INs (neurogliaform), and sncg-INs (CCK-basket cells) (Tasic et al., 2018). Our results show that VIP-INs provide only a small contribution to the enrichment in 5HT3AR-INs in PER (Figure 2 – Figure supplement 1). The other two groups lack specific mouse lines or unique molecular markers. We used instead in vitro electrophysiology to characterize the firing patterns of GABAergic interneurons in L5 of PER. We focused on L5 because it is where we found the largest difference in the percentage of GABAergic neurons not expressing PV and SST between PER and wS1 (Figure 1G-I). Neurogliaform cells in the cortex, hippocampus and amygdala show a “late-spiking” phenotype characterized by a long latency to the first spike, broad AHPs and no adaptation (Overstreet-Wadiche and McBain, 2015). Sncg-INs include CCK-basket cells, and show a regular firing pattern with little or no adaptation (Kawaguchi and Kubota, 1998). We obtained brain slices containing PER from GAD67-GFP mice and performed whole-cell patch clamp experiments in current clamp to measure voltage responses of GABAergic neurons. We found four types of firing patterns in PER (n= 33 cells): the fast-spiking phenotype (51.5%), associated to PV-INs, the irregular firing pattern (3.03%), associated to some VIP-INs, the late-spiking phenotype (39.4%), associated to neurogliaform cells, and the adapting firing pattern (6.07%), associated to some SST-INs. In L5 of wS1 we found five firing patterns (n= 20 cells): fast-spiking (55%), irregular (5%), late-spiking (10%), adapting (5%), quasi-fast spiking (25%), associated to layer 4 targeting non-Martinotti SST-INs (Ma et al., 2006; Nigro et al., 2018) (Figure 3A). Our electrophysiological characterization of the GABAergic population in PER and wS1 showed that late-spiking neurogliaform cells are enriched in L5 of PER. The late-spiking neurons in PER had short dendrites and a dense local axonal arborization, typical of neurogliaform cells in neocortical L1 and hippocampus (Overstreet-Wadiche and McBain, 2015) (Figure 3B). We conclude that neurogliaform cells might account for most of the difference in the density of 5HT3aR-INs between PER and wS1.

**Figure 3.**
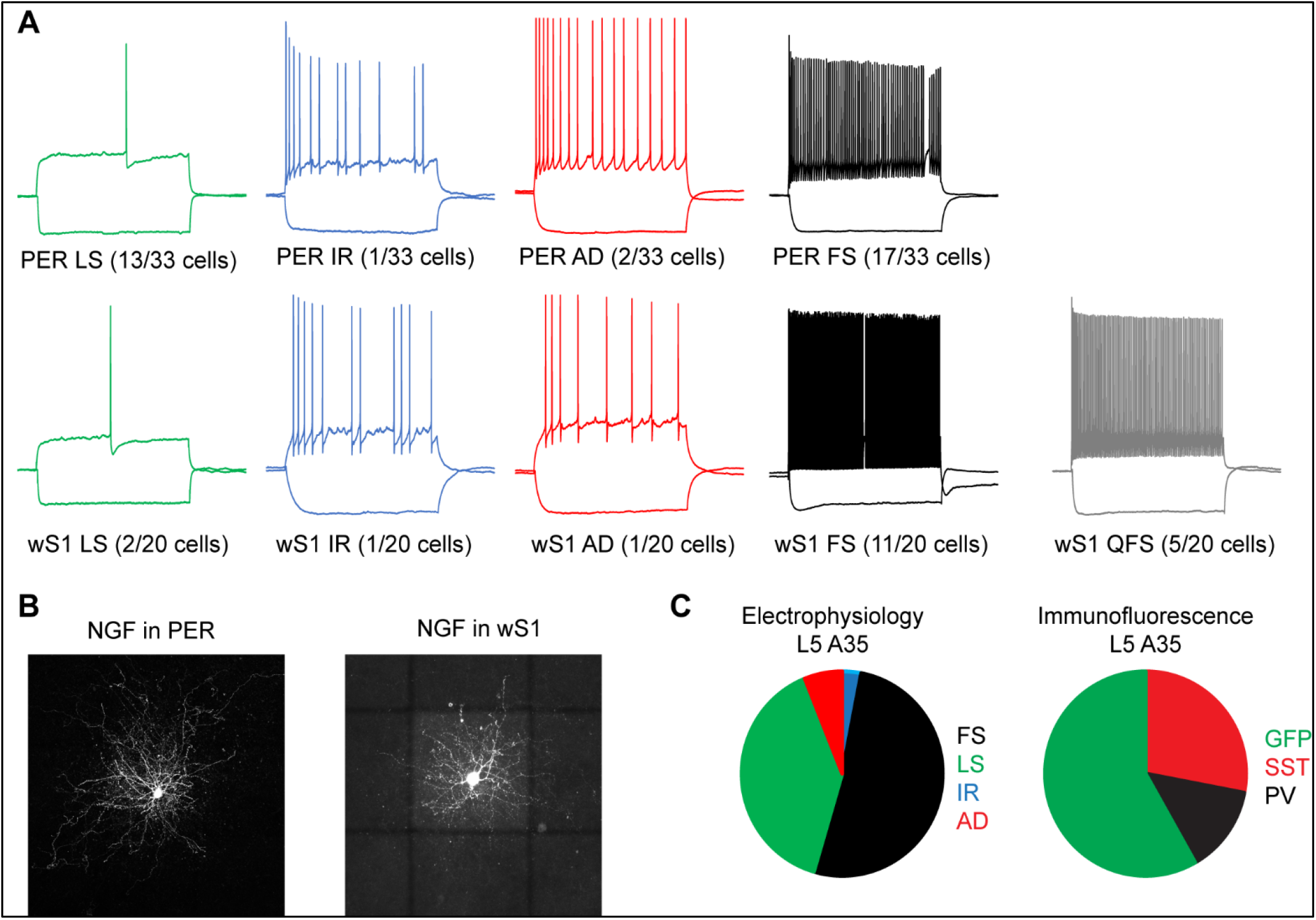
**A**. Firing patterns recorded from GABAergic neurons in L5 of A35 (upper row) and wS1 (lower row). **B**. Confocal stacks showing of a neurogliaform cell in PER (left) and wS1 (right). **C**. Comparison of the percentage of firing patterns (left) and molecularly defined populations (right) in L5 of A35. Note the high percentage of fast-spiking neurons as compared to the low percentage of PV-INs in PER.

### A fast-spiking population that do not express PV in PER

Unexpectedly, the fast-spiking phenotype accounted for the largest population in PER and wS1, which is at odds with the low density of PV-INs in PER (Figure 1 and Figure 3C). We hypothesized that some fast-spiking neurons in PER do not express detectable levels of PV. To test this hypothesis, we labelled fast-spiking neurons with a viral approach that exploits enhancer driven expression of tdTomato in PV-INs throughout the brain (Vormstein-Schneider et al., 2020). We injected the S5E2-dTom virus in PER of GAD67-EGFP mice and stained for SST to subtract the population of virus transfected neurons that co-expressed SST (Figure 4 – Figure supplement 1 and Figure 4 – Figure supplement 2). We found that the E2-dTom/SST-represented the largest GABAergic population in the deep layers of PER (Figure 4A-B). The percentage of E2-dTom/SST- was 47.1 ± 11.8% of the total GABAergic population in L5 of A35, which was very similar to the percentage of PV-INs in L5 wS1 (48.3 ± 8.3%) and to the percentage of fast-spiking neurons in L5 of PER (51.5%) (Figure 4C).

**Figure 4.**
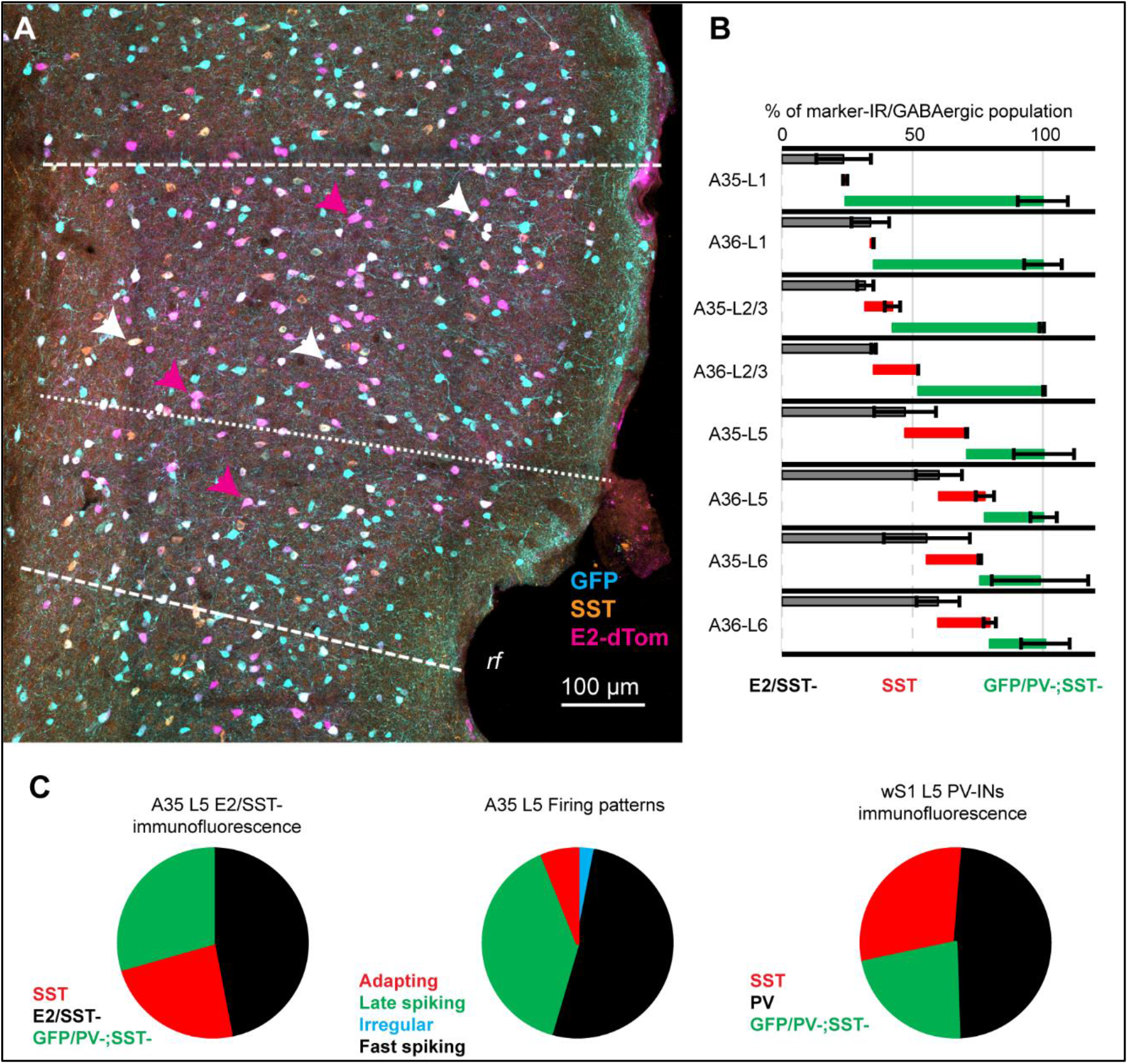
**A**. Representative confocal stack of immunofluorescence for GFP (turquoise) SST (orange) and dTomato (lilla) in PER of the GAD67-EGFP mouse line. **B**. Cumulative percentage of E2-dTom/SST- (black), SST-INs (red), GFP/PV-;SST- (green) neurons in the total GABAergic population of PER in the GAD67-EGFP mouse line (n= 2 mice). **C**. Pie charts comparing the percentage of marker defined populations in L5 A35 (left), electrophysiological characterization in L5 A35 (middle) and marker defined populations in L5 wS1. Note how the E2/SST-percentage is very similar to that of fast-spiking neurons in PER and PV-INs in wS1.

This result further suggests that PER is populated by a population of GABAergic neurons expressing a fast-spiking phenotype but not expressing PV. To directly test this hypothesis, we performed in vitro patch clamp recordings from brain slices of wild type C57 mice injected with the S5E2-dTom virus. We found that 88.9% (16 out of 18 neurons) of dTomato labelled neurons showed a fast-spiking phenotype and 11.2% showed an adapting (1/18) or low-threshold firing pattern (1/18), typical of SST neurons (Figure 5A-B, and Figure 5 – Figure supplement 1). We performed immunofluorescence for PV on 15 of the recorded fast-spiking neurons and found that 40% (6/15) did not express PV (E2-PV-) (Figure 5C). Our results show that the fast-spiking population of PER contains neurons that do not express PV, highlighting a novel diversity within this GABAergic type that is area specific.

**Figure 5.**
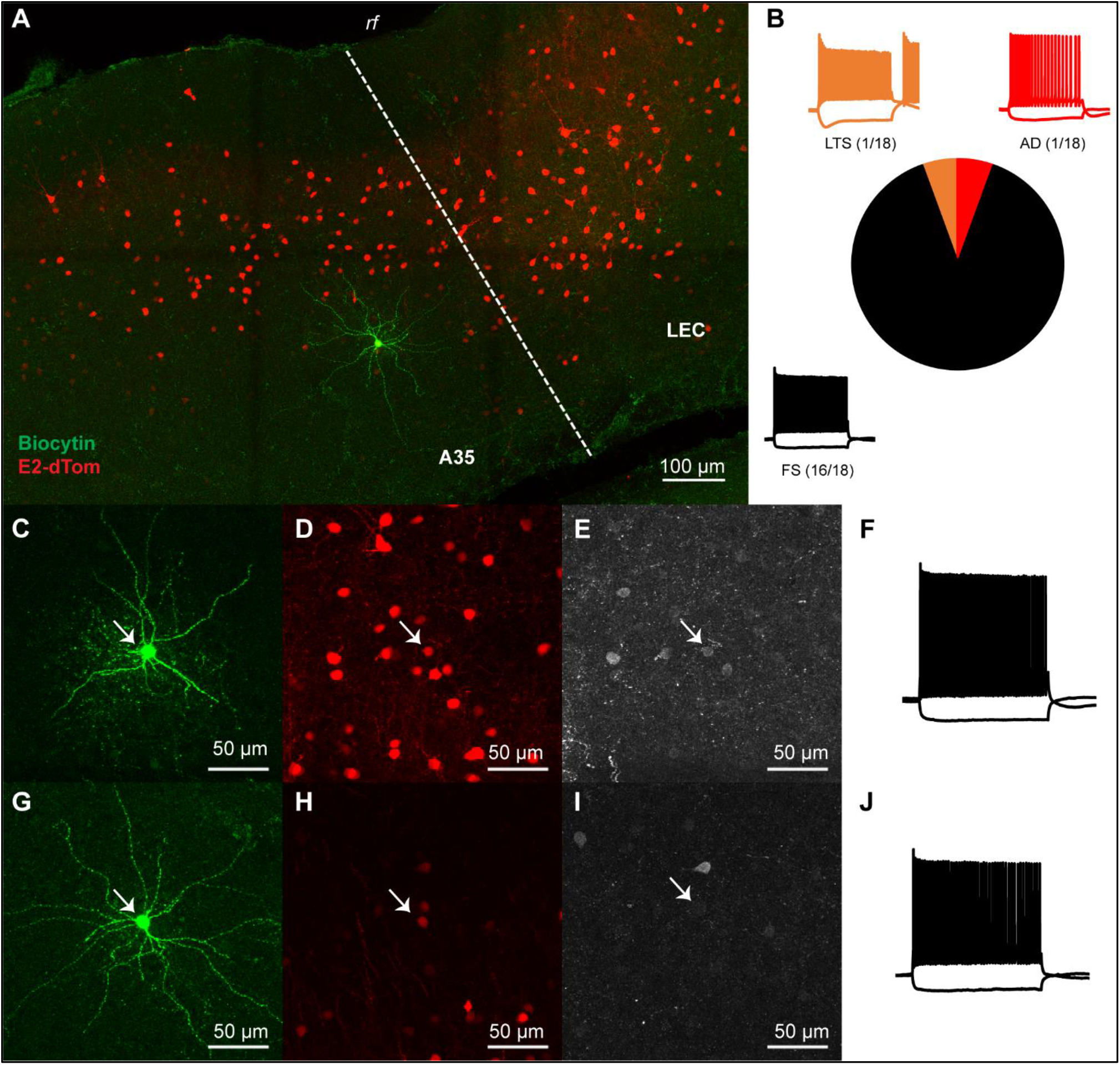
**A**. Confocal stack showing the location in L5 of A35 of a E2-dTom neuron filled with biocytin (green). Neurons labeled by the E2-dTom virus are shown in red. Dashed line shows the border with LEC. **B**. Pie chart showing the percent of LTS (orange), adapting (red) and fast-spiking (FS) (black) among E2-dTom neurons. **C-E**. Confocal stacks of a E2-dTom neuron (D, red) filled with biocytin (C, green) and expressing PV (E, white, white arrow). **F**. Voltage response of the cell shown in C-D to hyper- and depolarizing current steps. **G-I**. Confocal stacks of a E2-dTom neuron (H, red) filled with biocytin (G, green) and not expressing PV (I, white, white arrow). **J**. Voltage response of the cell shown in C-D to hyper- and depolarizing current steps.

### Fast-spiking interneurons in PER show different electrophysiological properties than in wS1

To test whether the firing patterns of fast-spiking neurons not-expressing PV in PER are different from PV-INs in PER and wS1, we performed a hierarchical cluster analysis on the electrophysiological properties (Figure 6A). The dendrogram shows that the population analyzed can be divided in two clusters, one containing all wS1 neurons, and a second containing all but 4 PER neurons (Figure 6A). The group containing perirhinal fast-spiking neurons included PV-INs and neurons not expressing PV, suggesting that these neurons show similar electrophysiological responses (Figure 6A). We further compared the electrophysiological parameters included in the cluster analysis (see Methods) and found that fast-spiking interneurons in PER had a significantly higher IR and as a consequence they fired action potentials in response to smaller current steps (Figure 6B and G). Fast-spiking neurons in PER had a less pronounced sag, which suggests a lower expression of HCN channels (Figure 6C). A hallmark of PV-INs are their fast APs and high firing rate, interestingly fast-spiking neurons in PER had wider APs, longer AHP and reached lower maximal firing rates than their wS1 counterparts (Figure 6D-G). We did not find any difference between E2-PV+ and E2-PV-neurons in PER, suggesting that these neurons might belong to the same type and differ only in expression of PV.

**Figure 6.**
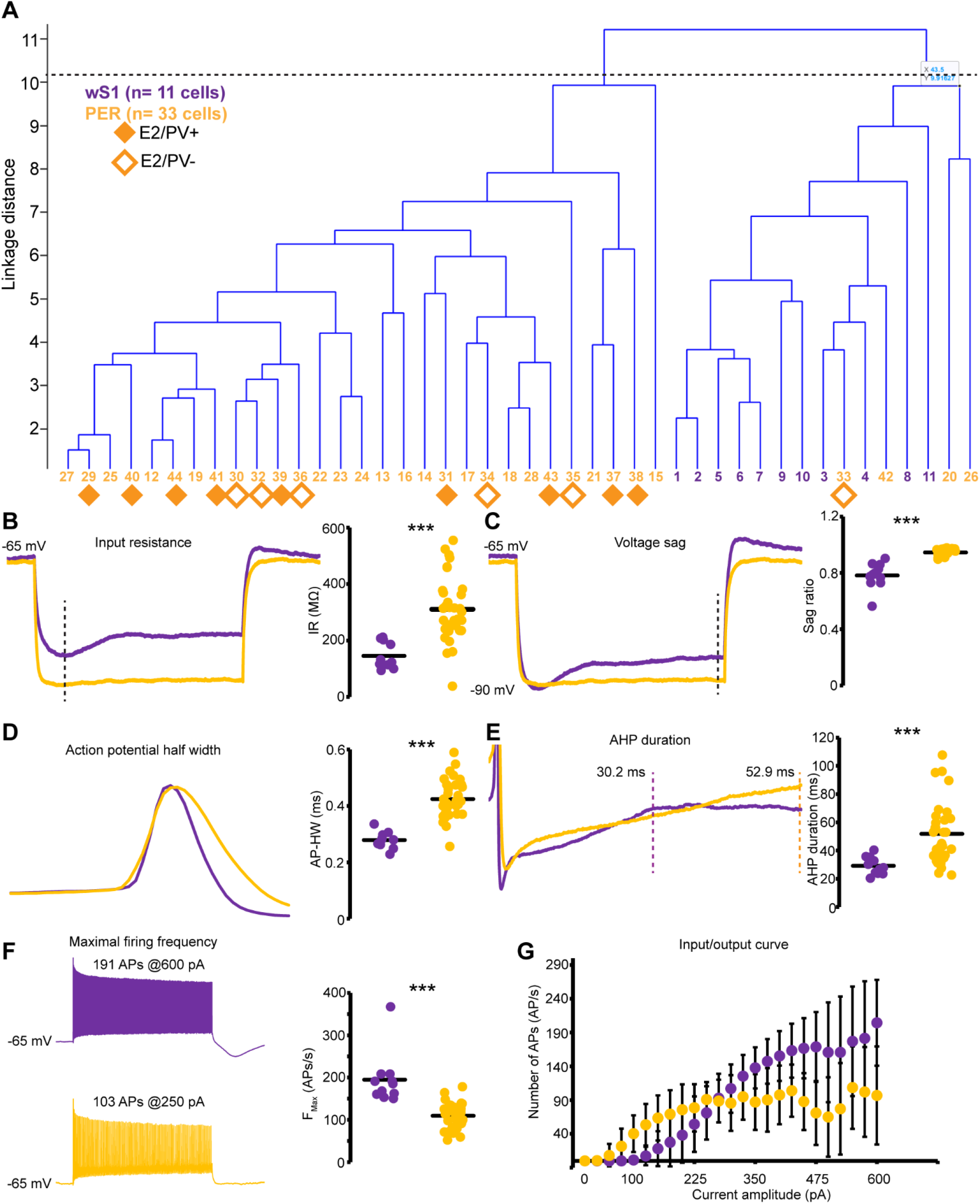
**A**. Dendrogram obtained from a hierarchical cluster analysis of 44 fast-spiking neurons (11 in wS1 and 33 in PER). The dashed line represents the level at which the clusters were identified by using the inconsistency index (see Methods). Numbers on the x-axis represent the cell’s IDs color-coded according to the cortical area (yellow, PER; purple, wS1). Filled diamonds indicate E2-dTom neurons expressing PV. Open diamonds indicate E2-dTom neurons not expressing PV. **B**. Representative voltage response of fast-spiking neurons to a -100 pA current step in wS1 (purple) in PER (yellow) neurons. The dashed line indicates the peak of the response. The plot on the right shows the IR of all neurons (circles) and the population mean (black line) (wS1, 145.4 ± 46.4 MΩ; PER, 310.7 ± 114.4 MΩ; p= 3.6E-8, T-test). **C**. Representative voltage response to a hyperpolarizing current step of fast-spiking neurons in wS1 (purple) and in PER (yellow). All responses used to calculate the sag ratio reached -90 ± 2 mV. Note the difference in the steady-state response (dashed line). The plot on the right shows the sag ratio of all neurons (circles) and the population average (black line) (wS1, 0.78 ± 0.1; PER 0.94 ± 0.02; p= 0.0001, T-test). **D**. Representative action potentials of fast-spiking neurons in wS1 (purple) and PER (yellow). The plot on the right shows the AP-HW of all neurons (circles) and the population mean (black line) (wS1, 0.28 ± 0.03 ms; PER, 0.42 ± 0.07 ms; p= 2.9E-12, T-test). **E**. Representative AHPs following single APs in wS1 (purple) and PER (yellow). The AHPdur is indicated by the two dashed lines. The plot on the right shows the AHPdur of all neurons (circles) and the population mean (black line) (wS1, 29.1 ± 6.2 ms; PER, 51.8 ± 21.3 ms; p= 2.4E-6, T-test). **F**. Representative responses of fast-spiking neurons in wS1 (purple) and PER (yellow) to depolarizing current steps eliciting maximal firing frequency. The plot on the right shows the Fmax in all neurons (circles) and the population mean (black line) (wS1, 194.8 ± 60.7 APs/s; PER, 109.5 ± 29.6 APs/s; p= 0.0008, T-test). **G**. Plot of the average number of APs evoked with current steps of increasing amplitude in wS1 (purple) and PER (yellow).

## Discussion

The present study offers a comprehensive description of the GABAergic diversity in the mouse PER, and demonstrates several differences compared to a sensory region (wS1). First, we demonstrate that PER contains a comparable density of GABAergic neurons as other cortical areas despite the low density of PV-INs. Second, PER is enriched in neurogliaform cells in layer 5. Since our molecular characterization showed no other candidate for the increased density of 5HT3aR-INs in other layers, we hypothesize that neurogliaform cells are more numerous throughout the layers of PER. Third, we corroborated previous observations of an increased SST-INs/PV-INs ratio in L2/3 (Kim et al., 2017). We extend this observation by showing that the density of SST-INs is significantly higher in PER as compared to wS1. Finally, we demonstrate that the fast-spiking population of PER is comparable to that of wS1 and most of these neurons do not express PV. We then show that the fast-spiking population of PER shows several electrophysiological differences with that in wS1 (and other telencephalic areas), highlighting a novel diversity within this electrophysiological cell-type.

### Enrichment of 5HT3aR-INs in PER

We found that, although the density of PV-INs is lower in PER than in wS1, the GABAergic population is of comparable size, or even larger (e.g., in L2/3 and 6). This difference is partially due to an enrichment of GABAergic neurons belonging to the neurogliaform type, and to a lesser extent to the VIP-expressing group. These GABAergic types belong to the CGE-derived class (Tremblay et al., 2016). Interestingly, transcriptomic analysis of the human temporal cortex showed that the human cortex is enriched of CGE-INs and intratelencephalic projecting excitatory cells (IT) (Hodge et al., 2019). CGE-INs and IT-projection neurons show a preferential connectivity that is established during development (Wester et al., 2019). The PER shows very sparse projections outside of the telencephalon, except for the thalamus, which suggests that a developmental program might be responsible for the enrichment of CGE-INs (McIntyre et al., 1996; Benavidez et al., 2021).

### Most fast-spiking GABAergic neurons in PER do not express PV

The low expression of PV in PER has been used by neuroanatomists to delineate PER for several decades in different species (Pitcänen and Amaral, 1993; Burwell et al., 1995; Uva et al., 2004; Beaudin et al., 2013). However, this observation has only recently been examined in the context of GABAergic cell-types and inhibitory circuits using specific mouse lines targeting PV-INs (Whissel et al., 2015; Kim et al., 2017). We have recently described that the labelling in the PV-IRES-Cre mouse line is very inefficient and quantifications of the number of PV-INs would be dramatically underestimated (Nigro et al., 2021). The present study corroborates the low density of PV-INs in PER by quantifying PV-IR neurons in PER and wS1. However, our electrophysiological characterization demonstrates that fast-spiking interneurons represent half of the GABAergic population in PER, similarly to wS1. Taking advantage of a newly developed viral approach we further demonstrate that most fast-spiking neurons in PER do not express PV. The electrophysiological properties of fast-spiking interneurons in PER differ from those in wS1. Our results suggest that the fast-spiking population of the cortex shows a yet unappreciated diversity. Recent transcriptomic examination of the GABAergic cell-types of the whole cortex did not identify transcriptomic types with signatures of fast-spiking neurons but not expressing PV (Yao et al., 2021). This is likely due to the use of specific mouse lines targeting different GABAergic types, which would prevent the identification of a population that do not express any of the main known GABAergic molecular markers.

### Ideas and hypotheses

The expression of PV is the hallmark of fast-spiking GABAergic neurons throughout the telencephalon (Yao et al., 2021). Our results raise the hypothesis that PV expression per se might not be enough to define this GABAergic class in all cortical areas. We have previously shown that the PV-IRES-Cre mouse line shows a low efficiency in several associative areas of the cortex (Nigro et al., 2021). The low density of PV-INs seems also to be a common feature of associative areas (Whissel et al., 2015; Kim et al., 2017). We hypothesize that our results might extend beyond PER to other associative areas of the cortex. Transcriptomic analysis of neurons labeled by the S5E2-dTom virus might reveal the taxonomic relationship between these different types of fast-spiking neurons in the cortex.

Until now we have assumed that fast-spiking neurons not expressing PV belong to the PV GABAergic class based on our electrophysiological characterization. The lack of PV expression might be due to plasticity of the circuit in which these cells are embedded. Indeed, PV expression is activity dependent and has been shown to be modulated by learning (Donato et al., 2013).

An alternative scenario could be that these neurons belong to a different class (e.g., Lamp5, scng, SST, or VIP), and simply lost their characteristic marker. Interestingly, the S5E2-dTom virus labels all PV-INs and about 60% of the SST population. The fast-spiking phenotype described here in perirhinal neurons is reminiscent of the quasi-fast spiking phenotype of non-Martinotti neurons described in L4 and L5b of wS1 (Ma et al., 2006; Nigro et al. 2018; Naka et al. 2019). However, our characterization of the firing pattern of SST neurons in L5 of PER shows that these neurons express either an adapting or a low-threshold spiking phenotype. Moreover, non-Martinotti neurons are extremely rare outside of wS1 (Ma et al., 2006; Scala et al., 2019). Finally, some chandelier cells do not express PV, and express a firing pattern similar to that described here in PER (Taniguchi et al., 2013). For these reasons we believe that the fast-spiking population in PER belongs to the PV transcriptomic class but lacks PV expression.

## Methods

### Animal models

The research described here was performed on adult mice (>8 weeks old) of both sexes. We used the following mouse lines: C57BL/6J line from Jackson Laboratories (stock 000664), GAD67-eGFP obtained as gift from Dr. Yanagawa (Tamamaki et al., 2003), the PV-IRES-Cre line (B6;129P2-Pvalbtm1(cre)arbr/J, stock 008069); R26R-EYFP (129×1-Gt(ROSA)26Sortm1(EYFP)Cos/J stock 006148) and the SST-Cre line (Ssttm2.1(cre)Zjh/J, stock 013044). We used two brains of the 5HT3aR-EGFP mouse line (MMRRC: Tg(Htr3a-EGFP)DH30Gsat/Mmnc stock: 000273-UNC) kindly donated by Dr. Ramesh Chittajallu. The PV-IRES-Cre line was crossed with the R26R-EYFP to obtain EYFP expression in PV-INs. The PV-IRES-Cre was bred as homozygous, whereas the GAD67-eGFP and the R26R-EYFP were bred as heterozygous. Animals were bred in house, group housed in enriched cages with water and food ad libitum. Animals were kept with an inverted dark/light cycle of 12h/12h.

### AAV2/1-S5E2-dTomato (S5E2-dTom) purification

The plasmid construct for the S5E2-dTom was purchased from Addgene (#135630) (Vormstein-Schneider et al., 2020). We packaged the construct in AAV 2/1 serotype (a mosaic of capsid 1 and 2) and generated injection-grade viral vector using Heparin column affinity purification method (Nair et al iScience 2020). All the plasmids for AAV vector preparations were made using endotoxin free plasmid maxiprep kit (#12663, Qiagen). AAV 293 cells (#CVCL_6871, Agilent, USA) used for vector productions were maintained in DMEM (# 41965062, Thermo Fisher Scientific) containing 10% fetal bovine serum (#16000-044, ThermoFisher, USA) and penicillin/streptomycin antibiotics (#15140122, Thermo Fisher Scientific). The day before transfection, we seeded approximately 7x 106 AAV 293 cells on 150 mm cell culture plates and maintained at 37°C, 5% CO2. For the calcium chloride mediated co-transfection we used 22.5 μg pAAV-containing the transgene, 22.5 μg pHelper (#240071, Agilent, USA), 11.3 μg pRC (#240071, Agilent, USA) and 11.3 μg pXR1 (NGVB, IU, USA) capsid plasmids. The culture medium was replaced after 7 hours with fresh 10% FBS containing DMEM. Transfected cells were scrapped out after 72 hours and centrifuged at 200xg. The pellet was subjected to lysis using a buffer containing 150 mM NaCl-20 mM Tris pH 8.0 and 10% sodium deoxy cholate. The lysate was then treated with Benzonase nuclease HC (#71206-3, Millipore) for 45 minutes at 37 oC. Subsequently, the benzonase treated lysate was centrifuged at 3000xg for 15 mins. The clear supernatant was then subjected to HiTrap® Heparin High Performance (#17-0406-01, GE) affinity column chromatography using a peristaltic pump. The elute from the Heparin column was then concentrated using Amicon Ultra centrifugal filters (#Z648043, Millipore). We performed quantitative PCR on the viral stock and the titer was determined as approximately 1011 infectious particles/ml.

### Injections of S5E2-dTomato viral vector

The AAVs were injected into the brains of C57 (n= 6), GAD67-EGFP (n= 2) and on PV-EYFP (n= 2) mice. Mice were anaesthetized with isoflurane (4% Nycomed, airflow 1 l/min) in an induction chamber. The mice were placed in a stereotactic frame on a heated pad (37 °C) throughout the procedure and head-fixed with ear bars. Eye ointment was applied to protect the cornea. The following analgesic were applied subcutaneously: buprenorphine hydrochloride (0.1 mg/Kg, Temgesic, Invidior), meloxicam 1 mg/Kg, Metacam Boerringer Ingelheim Vetmedica), bupivacaine hydrochloride (1 mg/Kg locally, Marcain, Astra Zeneca). The skin on the incision site was shaved and disinfected with Pyrisept. An incision was made to access the skull. The skull was thinned with a dental drill at the desired location, and a hole was punched with the tip of a glass pipette. We used the following coordinates from bregma: for wS1 -0.7 mm AP, +3.1 mm ML, -0.15 and 0.55 DV; for PER -3 mm AP, -1.5 DV, for the ML the pipette was moved to the edge of the skull and then retracted 0.4 mm. Injections of 100-200 nl (50 nl/minute) in each location were made with a glass pipette (20-30 μm tip size) attached to an injector (Nanoliter 2010, World Precision Instruments) controlled by a microsyringe pump controller (Micro4 pump, World Precision Instruments). After retracting the pipette, the wound was rinsed with saline and sutured. The mice were allowed to recover in a heated chamber (37 °C) before returning to the homecage. Postoperative analgesic was applied 6-8 h after the procedure (Temgesic) and 24 h after the procedure (Metacam). The survival time for transduction and expression of the genetic material was 10-15 days.

### In vitro electrophysiology

Mice of either sex (>8-week-old) were euthanized with an overdose of pentobarbital (i.p. 100 mg/Kg, Apotekerforeninger), before intracardial perfusion with cutting solution (RT) of the following composition (in mM): 93 choline chloride, 3 KCl, 1.25 NaH2PO4, 30 NaHCO3, 20 HEPES, 10 glucose, 5 MgCl2, 0.5 CaCl2, 5 N-acetylcysteine, saturated with 95% O2 and 5% CO2. The brain was extracted and a section containing either PER or wS1 was glued onto the stage of a vibratome (VT1000, Leica) filled with cold (4°C) cutting solution. Slices (300 μm) were transferred to a chamber filled with warm (34°C) cutting solution and incubated for 15 minutes. The slices were stored in a different chamber filled with the following solution (in mM) (RT): 92 NaCl, 3 KCl, 1.25 NaH2PO4, 30 NaHCO3, 20 HEPES, 10 glucose, 5 MgCl2, 0.5 CaCl2, 5 N-acetylcysteine, saturated with 95% O2 and 5% CO2.

The slices were placed in a recording chamber of an upright microscope (BW51, Olympus) filled with warm (35°C) recording solution of the following composition: 124 NaCl, 3 KCl, 1.25 NaH2PO4, 26 NaHCO3, 10 glucose, 1 MgCl2, 1.6 CaCl2, saturated with 95% O2 and 5% CO2. Characterization of the electrophysiological properties was performed in presence of synaptic blockers: 10 μM DNQXNa2, 25 μM APV and 10 μM gabazine. wS1 was identified under DIC optics by the presence of characteristic barrels in layer 4. To identify A35 under DIC optics, we first identified the lateral entorhinal cortex (LEC) by its characteristic large neurons in layer 2a, and by the clear separation between layer 2a and 2b. A35 was defined as the cortex dorsally located to LEC. Only healthy, fluorophore-expressing neurons were selected for recordings and no other criterion was applied. Whole-cell patch clamp recordings were performed with borosilicate pipettes (Sutter Instruments) with a resistance of 3-7 MΩ filled with the following solution (in mM): 130 K-gluconate, 10 KCl, 10 HEPES, 0.2 EGTA, 4 ATP-Mg, 0.3 GTP-Na, 5 phosphocreatine-Na2, pH 7.3. In all recordings biocytin was added to the pipette solution to recover the layer location of the neuron. Membrane potentials reported were not corrected for a junction potential of -14 mV. Current clamp recordings were performed with a Multiclamp 700B amplifier (Molecular Devices), digitized with a 1550A Digidata (Molecular Devices), interfaced with a personal computer with pClamp 11 (Molecular Devices). Data were sampled at 40 kHz and low-passed filtered at 10 kHz. All recordings were performed from a holding potential of -65 mV. The electrophysiological parameters analyzed were defined as follows:

Input resistance (IR, MΩ): resistance measured from Ohm’s law from the peak of voltage responses to -25 pA hyperpolarizing current steps.

Sag ratio (dimensionless): measured from voltage responses peaking at -90 ± 2 mV to hyperpolarizing current steps. It is defined as the ratio between the voltage at the steady-state response and the peak.

Action potential threshold (APthr, mV): measured from action potentials (AP) evoked near rheobase, and defined as the voltage where the rise of the AP was 20 mV/ms.

AP half width (HW, ms): duration of the AP at half amplitude from APthr, measured from action potentials (AP) evoked near rheobase.

Amplitude of the fast afterhyperpolarization (fAHP, mV): measured from APthr, from action potentials (AP) evoked near rheobase.

AHP duration (AHPdur, ms): measure from action potentials (AP) evoked near rheobase, and defined as the time difference between the fAHP and the most depolarized voltage in a 200 ms window.

Maximal firing frequency (Fmax, AP/s): maximal firing frequency evoked with 1-s-long depolarizing current steps.

Rheobase (Rheo, pA): current level that evoked the first spike during a 1-s-long depolarizing ramp (500 pA/s).

### Histology

Mice used for histological analysis were anaesthetized with isoflurane and then euthanized with an overdose of pentobarbital (i.p. 100 mg/Kg, Apotekerforeninger). The mice were intracardially perfused with Ringer solution (0.85% NaCl, 0.025% KCl, 0.02% NaHCO3) followed by 4% parafolmaldehyde (PFA) in phosphate buffer (PB) (pH 7.4). The brain was extracted from the skull and placed in 4% PFA for 3 hours before moving it to a solution containing 15% sucrose in PB overnight. The brains were then stored in 30% sucrose for two days before being sliced at 50 μm with a freezing microtome. Slices were collected in six equally spaced series and placed in tubes containing a solution composed of 30% glycerol, 30% ethylene glycol and 40% phosphate buffer saline (PBS). The tissue was stored at -20 °C until used for histology.

The slices were washed in PB (3 × 10 minutes) before incubation with blocking solution (0.1% Triton-X, 10% NGS in PB). After blocking the tissue was incubated with the primary antibody for three days at 4 °C in a solution containing: 0.1% Triton-X, 1% NGS in PB. After being washed (3 × 1 hour in PB), the tissue was incubated with the secondary antibody overnight at 4 °C. The slices were washed (3 × 10 minutes in PB) and mounted on SuperFrost slides (Termo Fisher Scientific) and left to dry overnight. The slides were coversliped with entellan xylene (Merck Chemicals, Darmstadt, Germany) after being washed in xylene for 15 minutes, or with Fluoromount after being washed in distilled water for about 30 seconds. We used the following primary antibodies: Guinea Pig IgG anti-NeuN (1:1000, Sigma Millipore, #ABN90P), Rabbit anti-parvalbumin (1:1000, Swant, #PV-27), Mouse IgG1 anti-parvalbumin (1:1000, Sigma, #P3088), Rat IgG2a anti-RFP (1:1000, Proteintech, #5f8), Chicken IgY anti-GFP (1:1000, Abcam, #ab13970), Rabbit IgG anti-SST (1:1000, BMA Biomedicals), Rabbit IgG anti-VIP (1:1000, Immunostar). We used the following secondary antibodies: Goat anti-guinea pig (IgG H+L) A647 (1:500, Invitrogen, #A-21450), Goat anti-Rat (IgG H+L) A-546 (1:500, Invitrogen, #A11081), Goat anti-Rabbit (IgG H+L) A488 (1:500, Invitrogen, #A11008), Goat anti-chicken (IgY H+L) A488 (1:500, Invitrogen, #A11039), Goat anti-rabbit (IgG H+L) A546 (1:500, Invitrogen, #A11010), Goat anti-rabbit (IgG H+L) A635 (1:500, Invitrogen, A31576).

### Image acquisition and analysis

We used a confocal microscope (Zeiss LSM 880 AxioImager Z2) to image regions of interest (ROIs) to count labeled neurons. Images of PER and wS1 were taken with a x20 air objective with 1 Airy unit pinhole size and saved as czi files. We selected 3 to 5 ROIs for each cortical region. ROIs contained the whole cortical area of interest in a given slice. Files containing the ROIs were uploaded in Neurolucida (Micro Bright Field Bioscience) for analysis. We created contours to delineate the layers of the ROIs using the NeuN signal. The PER was divided in area 35 (A35) and area 36 (A36), and delineated according to Beaudin et al. (2013). Symbols for each signal were used to count cells labeled by different markers. We counted all cells contained within the contours. Since we counted slices distanced 300 μm from each other, overcounting in the z axis is not relevant and we did not apply any correction. Quantification of the number of markers in each contour was done in Neurolucida Explorer (Micro Bright Field Bioscience) and exported in excel files. The density of neurons labeled by a marker was measured as the number of labeled neurons in a contour divided by the area of the contour (cells/mm2). The GAD67-eGFP mouse line labelled virtually all PV-INs and VIP-INs, but showed a lower specificity for SST-INs (Figure 1 – Figure supplement 1). For this reason, the number of PV-INs, SST-INs and VIP-INs was measured from the immunolabelling for the specific marker, independently of GFP expression. Transcriptomic analysis of the neuronal population of the cortex shows that SST, VIP and PV are expressed exclusively in GABAergic neurons. We excluded from the counting a small population of PV-IR pyramidal neurons in L5B of wS1 (van Brederode et al., 1991). Values are reported as mean ± standard deviation). All graphs were created in excel, images of ROIs were created in Zen lite (Zeiss) and figures in illustrator.

### Statistics

Principal component analysis on the densities of different GABAergic neurons across layers of PER and wS1 was performed in Matlab 2021 (Mathworks) using the function “pca”. We obtained the explained variance for each principal component, the principal component coefficients, the principal component scores, and the principal component variances. Hierarchical cluster analysis was performed in Matlab 2021 (Mathworks). After normalization with the function “zscore”, we calculated the distance between pairs of objects with the function “pdist” and the method “cityblock”. Objects were grouped according to the calculated distances with the function “linkage” and the method “average”. After calculating the inconsistency index (“inconsistent”), clusters were obtained by using the function “cluster” and the inconsistency index as cutoff. This method identified two clusters.

We used one-way ANOVA with Bonferroni correction for statistical comparison of different layers of A35, A36 and wS1. We used the number of animals (n= 5) as sample population in all comparisons but for the 5HT3AR-EGFP neurons, where the comparison was performed on slices (n= 10), because the number of available brains was too low (n= 2). We did not use any statistical method to pre-determine sample size. The number of mice, slices or neurons used for statistical comparison are consistent with other studies in the field (Ma et al., 2006; Whissel et al., 2015; Kim et al., 2017; Nigro et al., 2018). No blinding method was used.

## Acknowledgments

We thank Dr. Ramesh Chittajallu for the kind donation of the brains of the 5HT3aR-EGFP mice and Dr. Timothy Petros for helping with obtaining the brain of the 5HT3aR-EGFP mice.

---

**Figure 1 – Figure supplement 1.**
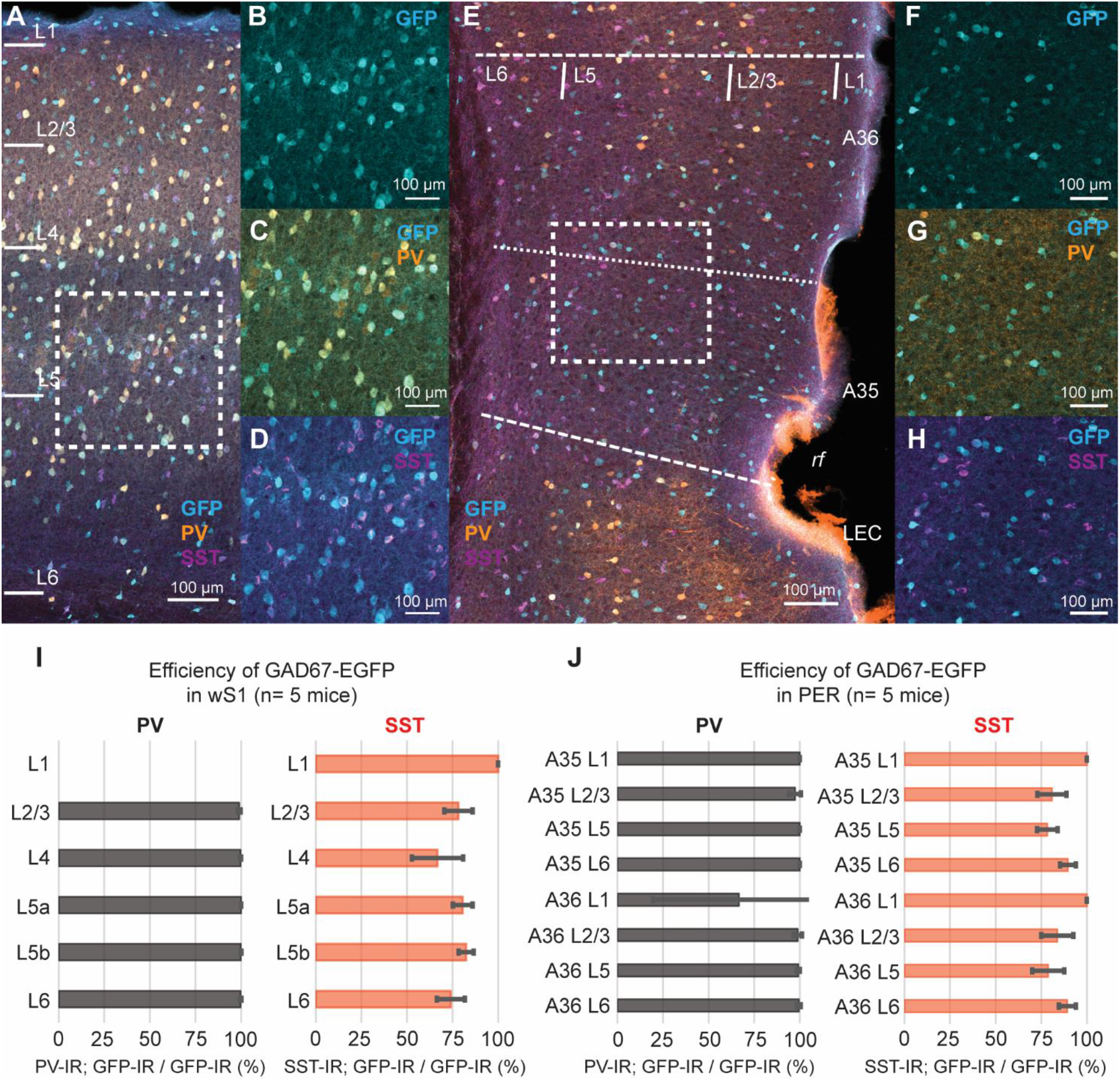
**A**. Representative confocal stack of immunofluorescence for GFP (turquoise), PV (orange) and SST (purple) in the wS1 of a GAD67-EGFP mouse. **B-D**. Magnification of the area in the dotted rectangle in A. **E**. Representative confocal stack of immunofluorescence for GFP (turquoise), PV (orange) and SST (purple) in PER of a GAD67-EGFP mouse. **F-H**. Magnification of the area in the dotted rectangle in E. **I**. Bar plots showing the efficiency of the GAD67-EGFP mouse line for PV (black, left) and SST (red, right) in wS1. **J**. Bar plots showing the efficiency of thee GAD67-EGFP mouse line for PV (black, left) and SST (red, right) in PER.

**Figure 1 – Figure supplement 2.**
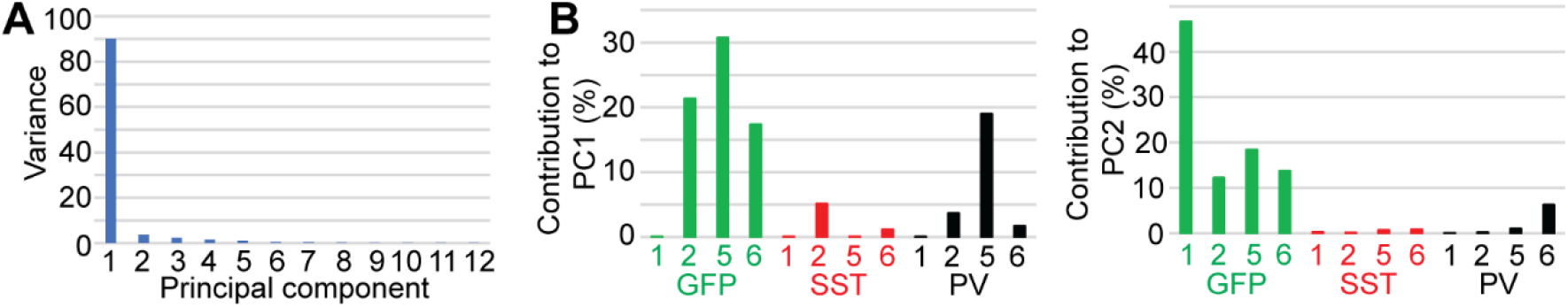
**A**. Bar plot showing the variance explained by each principal component. **B**. Bar plots showing the contribution of each variable (i.e., marker density in a given layer) to the first (left) and second (right) principal component.

**Figure 1 – Figure supplement 3.**
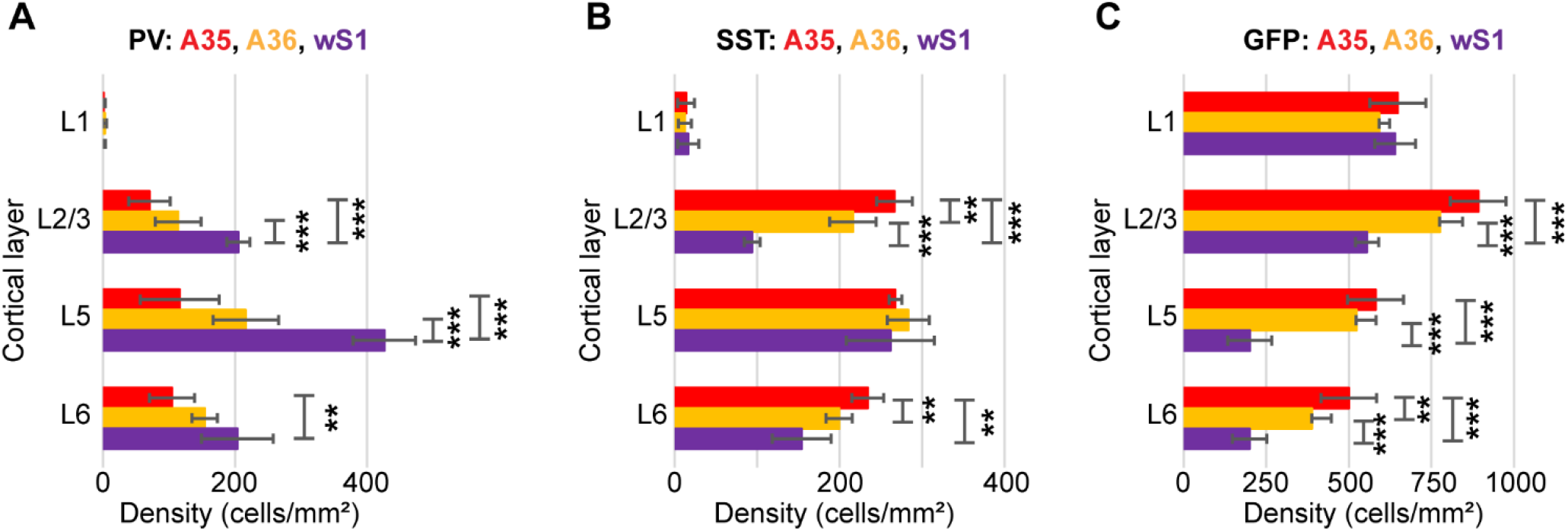
**A**. Bar plot showing the statistical comparison of the density of PV-IR neurons in the GAD67-EGFP mouse line (n= 5 mice): L1 (F= 2.06, df= 2, p= 0.17, One-way ANOVA); L2/3 (F= 28.54, df= 2, p= 2.75E-5, One-way ANOVA; A35 vs wS1, p= 3.16E-5, Bonferroni correction; A36 vs wS1= 0.0007, Bonferroni correction); L5 (F= 45.5, df= 2, p= 2.5E-6, One-way ANOVA; A35 vs wS1, p= 2.37E-5, Bonferroni correction; A36 vs wS1, p= 0.0001, Bonferroni correction); L6 (F= 8.17, df= 2, p= 0.006, One-way ANOVA; A35 vs wS1, p= 0.011, Bonferroni correction). **B**. Bar plot showing the statistical comparison of the density of SST-IR neurons in the GAD67-EGFP mouse line: L1 (F= 0.21, df= 2, p= 0.82, One-way ANOVA); L2/3 (F= 87.06, df= 2, p= 7.18E-8, One-way ANOVA; A35 vs A36, p= 0.014, Bonferroni correction; A35 vs wS1, p= 6.85E-6, Bonferroni correction; A36 vs wS1, p= 0.0003, Bonferroni correction); L5 (F= 0.53, df= 2, p= 0.6, One-way ANOVA); L6 (F=12.83, df= 2, p= 0.001, One-way ANOVA; A35 vs A36, p= 0.015, Bonferroni correction; A35 vs wS1, p= 0.004, Bonferroni correction). **C**. Bar plot showing the statistical comparison of the density of GFP-IR/-PV-SST neurons in the GAD67-EGFP mouse line: L1 (F= 1.61, df= 2, p= 0.24, One-way ANOVA); L2/3 (F= 51.43, df= 2, p= 1.3E-6, One-way ANOVA; A35 vs wS1, p= 6.04E-6, Bonferroni correction; A36 vs wS1, p= 0.0007, Bonferroni correction); L5 (F= 56.88, df= 2, p= 7.55E-7, One-way ANOVA; A35 vs wS1, p= 1.21E-5, Bonferroni correction; A36 vs wS1, p= 4.16E-5, Bonferroni correction); L6 (F= 38.09, df= 2, p= 6.35E-6, One-way ANOVA; A35 vs A36, p= 0.014, Bonferroni correction; A35 vs wS1, p= 1.99E-5, Bonferroni correction; A36 vs wS1, p= 0.0007, Bonferroni correction).

**Figure 2 – Figure 2 supplement 1.**
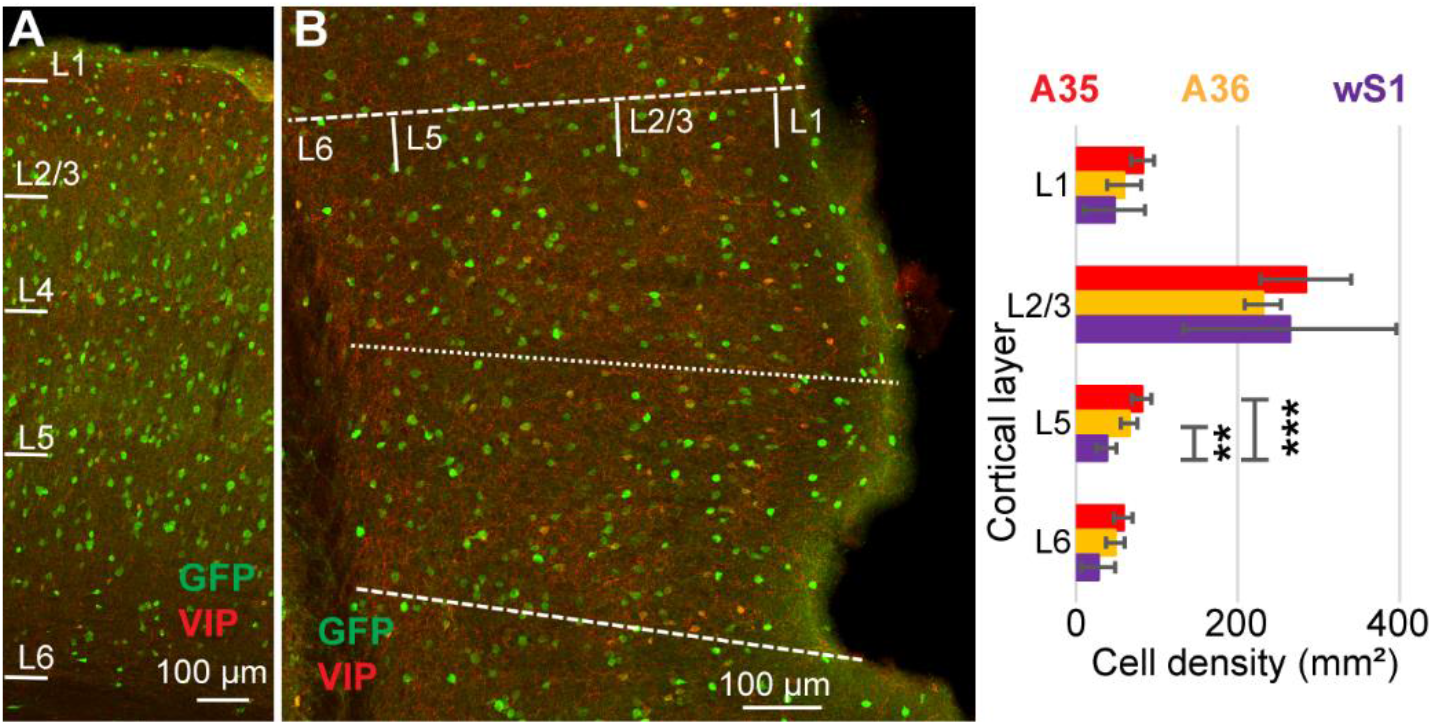
**A**. Representative confocal stack of immunofluorescence for GFP (green) and VIP (red) in wS1 of the GAD67-EGFP mouse line. **B**. Representative confocal stack of immunofluorescence for GFP (green) and VIP (red) in PER of the GAD-EGFP mouse line. **C**. Bar plot showing the statistical comparison of the density of VIP-IR neurons across layers of A35 (red), A36 (yellow) and wS1 (purple) (n= 5 mice): L1 (F= 2.34, df= 2, p= 0.14, One-way ANOVA); L2/3 (F= 0.51, df= 2, p= 0.61, One-way ANOVA); L5 (F= 17.76, df= 2, p= 0.0003, One-way ANOVA; A35 vs wS1, p= 0.0005, Bonferroni correction; A36 vs wS1, p= 0.005, Bonferroni correction); L6 (F= 5.22, df= 2, p= 0.02, One-way ANOVA).

**Figure 4 – Figure supplement 1.**
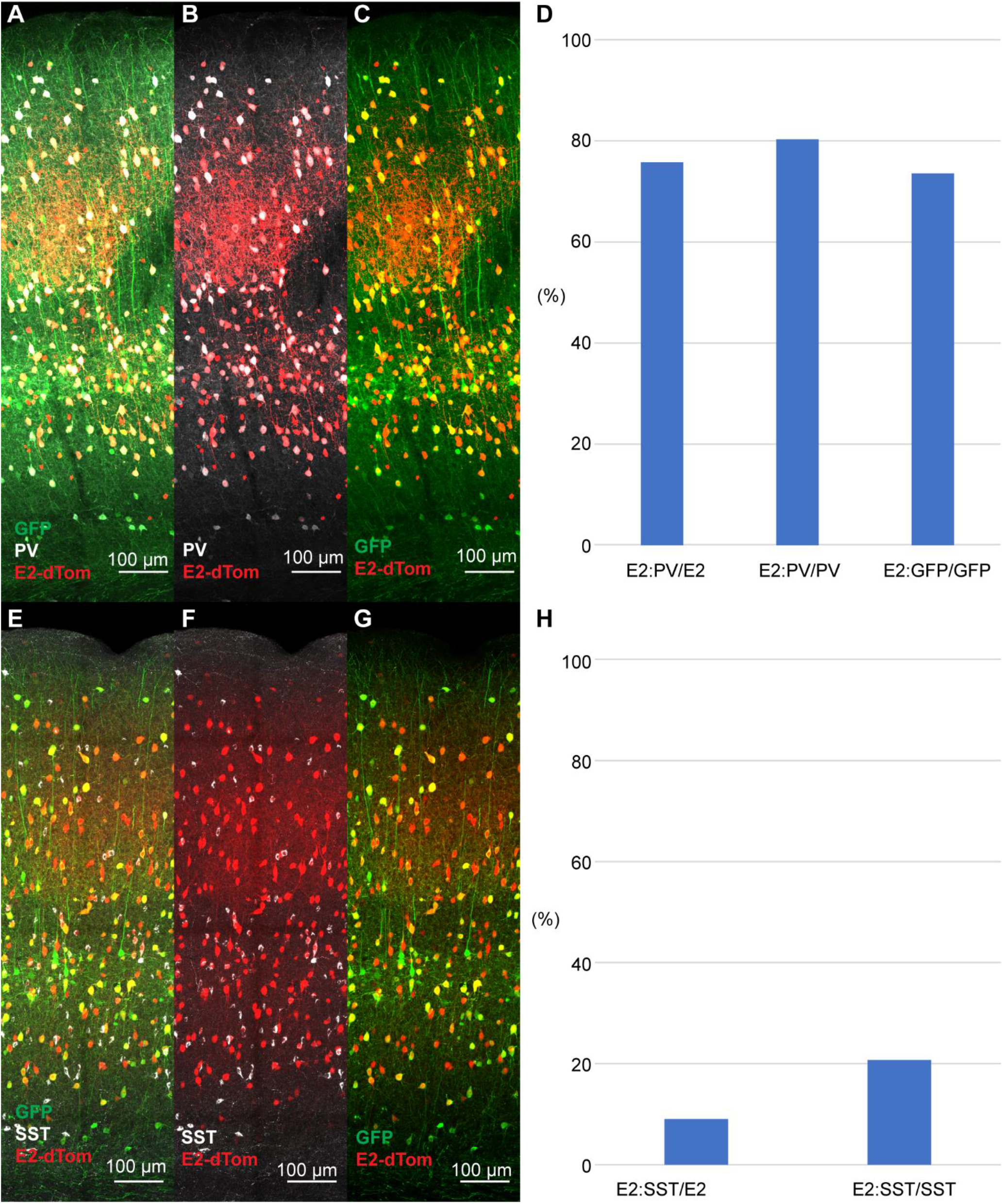
**A-C**. Representative confocal stacks of immunofluorescence for GFP (green), PV (white) and dTom (red) in a PV-IRES-Cre mouse injected with the S5E2-dTom virus in wS1. **D**. Bar plot showing the specificity (E2:PV/E2), and efficiency (E2:PV/PV and E2:GFP/GFP) of the S5E2-dTom virus in wS1 (n= 1 mouse). **E-G**. Representative confocal stacks of immunofluorescence for GFP (green), SST (white) and dTom (red) in PV-IRES-Cre mice injected with the S5E2-dTom virus in wS1. **H**. Bar plot showing the percent of neurons expressing dTomato (E2) that co-express SST (E2:SST/E2), and the percent of SST neurons labeled by the S5E2-dTom virus (E2:SST/SST) (n= 1 mouse).

**Figure 4 – Figure supplement 2.**
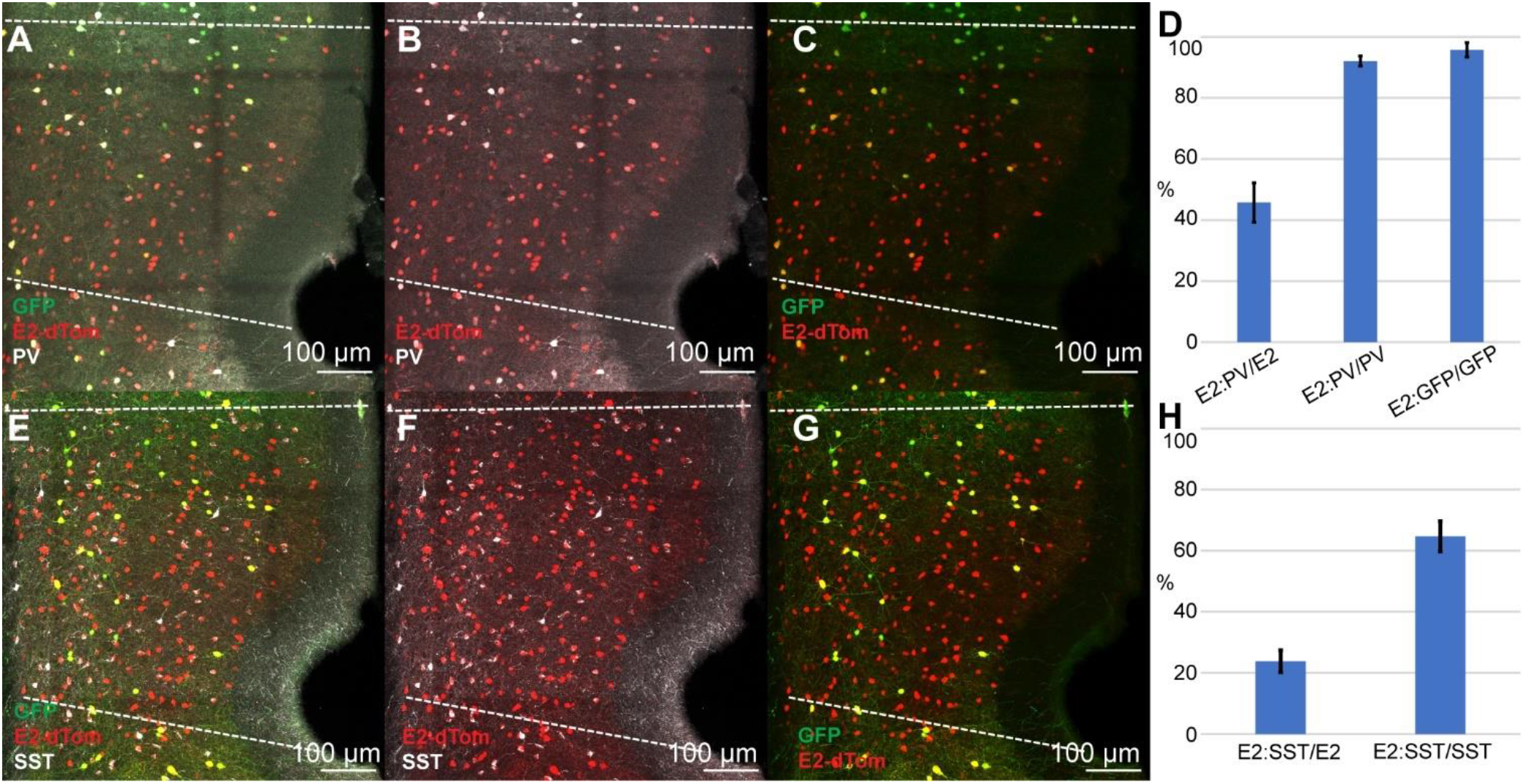
**A-C**. Representative confocal stacks of immunofluorescence for GFP (green), E2-dTom (red) and PV (white) in PV-IRES-Cre mice injected with the S5E2-dTom virus in PER. **D**. Bar plot showing the specificity (E2:PV/E2), and efficiency (E2:PV/PV and E2:GFP/GFP) of the S5E2-dTom virus in PER (n= 2 mice). **E-G**. Representative confocal stacks of immunofluorescence for GFP (green), SST (white) and dTom (red) in PV-IRES-Cre mice injected with the S5E2-dTom virus in PER. **H**. Bar plot showing the percent of neurons expressing dTomato (E2) that co-express SST (E2:SST/E2), and the percent of SST neurons labeled by the S5E2-dTom virus (E2:SST/SST) (n= 2 mice).

**Figure 5 – Figure supplement 1.**
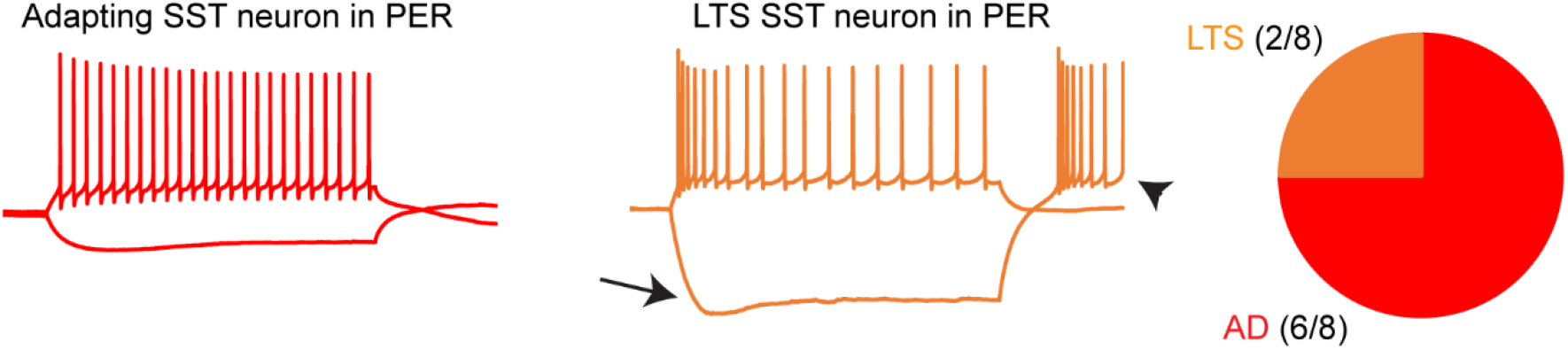
Representative voltage responses of an adapting (red) and a LTS (orange) neurons recorded in the SST-Cre mouse line and labeled with tdTomato. The pie chart shows the percentage of adapting (red) and LTS (orange) neurons in the SST-Cre line.

